# The mammalian decidual cell evolved from a cellular stress response

**DOI:** 10.1101/246397

**Authors:** Eric M. Erkenbrack, Jamie D. Maziarz, Oliver W. Griffith, Cong Liang, Arun R. Chavan, Günter P. Wagner

## Abstract

Among animal species, cell types vary greatly in terms of number and kind. The broad range of number of cell types among species suggests that cell type origination is a significant source of evolutionary novelty. The molecular mechanisms giving rise to novel cell types, however, are poorly understood. Here we show that a novel cell type of eutherian mammals, the decidual stromal cell (DSC), evolved by rewiring an ancestral cellular stress response. We isolated the precursor cell type of DSCs, endometrial stromal fibroblasts (ESFs), from the opossum *Monodelphis domestica*. We show that, in opossum ESF, the majority of decidual core regulatory genes respond to decidualizing signals, but do not regulate decidual effector genes. Rather, in opossum ESF, decidual transcription factors function in apoptotic and oxidative stress response. We propose that the rewiring of cellular stress responses could be a general mechanism for the evolution of novel cell types.

---

Multicellular organisms consist of numerous specialized cells, or cell types, that are produced *de novo* during development in each generation and that perform diverse physiological and structural functions. In evolution, sister cell types originate by differentiation from an ancestral cell type by a process called cell type splitting^1,2^. According to this model, novel cell types have arisen from ancestral cell types through modification of developmental programs leading to two derived cell types. The evolutionary diversification of cell types in metazoans has been a significant source of evolutionary novelty and was essential to the elaboration of increasingly complex body plans. While it is clear that cell types have diversified prodigiously in evolution, the molecular mechanisms leading to the origination of a novel cell type are not well understood^3^.

The evolution of mammalian pregnancy offers an opportunity to investigate cell type origination. Intensive selective pressures during the evolution of mammalian pregnancy led to the evolution of many functional specializations of the uterus that accommodate the implantation of the embryo and development of the placenta^4^. These novelties include the origin of specialized cell types such as the decidual stromal cell (DSC), the uterine natural killer cell (uNK), and a specialized form of resident macrophages^5^.

During both the menstrual cycle and pregnancy, DSCs differentiate from endometrial stromal fibroblasts (ESFs) on exposure to progesterone and signals from the embryo^6^. Responding to these signals and driving DSC differentiation is a complex gene regulatory network (GRN). Numerous transcription factors have been shown to transcriptionally and post-translationally interact to regulate effector gene sets conferring DSC cell type identity. Evolutionarily, phylogenetic cell type studies make clear that eutherian ESF and DSC are sister cell types^7^, and while ESFs are found in the oviduct of numerous amniotes, DSCs are exclusive to eutherians^8^. Moreover, it is clear that DSCs evolved from an ancestral ESF cell type, hereafter referred to as paleo-ESF, that existed in the phylogenetically recent stem lineage of eutherian mammals, having diverged 65 to 80 Mio years ago^9^, *i.e.* after the most recent common ancestor of marsupials and eutherians^10^. Hence, the evolutionary origin of DSC is an outstanding model to investigate the molecular mechanisms that led to the origin of a novel cell type.

To characterize the origin of the decidual stromal cell type, we isolated ESFs of the marsupial grey short tailed opossum *Monodelphis domestica,* hereafter called MdESF, which we use as a proxy for paleo-ESF. In humans and other eutherians, neo-ESF differentiates into DSC *in utero* when exposed to progesterone (P4) and estrogen, as well as ligands upstream of cyclic AMP (cAMP)/PKA signaling such as PGE2^11-13^, RLN^14^, or CGB (hCG)^15^. We assayed the response of MdESF to these stimuli that differentiate HsESF to HsDSC *in vitro* in order to identify the ancestral gene regulatory program from which the core network of DSC evolved. We find, surprisingly, that core components of the decidual GRN are responsive to progesterone and cAMP in opossum ESF, but rather than undergoing DSC differentiation, these cells undergo a cellular stress response.

## ESF isolation from *Monodelphis domestica* (40/40)

We utilized an established protocol to isolate MdESF by Percoll column gradient^16^. We validated by immunostaining and Western-blotting that cells isolated by this procedure are positive for the mesenchymal marker vimentin and negative for the epithelial marker cytokeratin (Supplementary Information Fig. 1a). Relative to other layers in the column, these cells expressed higher levels of the ESF markers *HOXA11*, *HOXA10,* and *PGR*. We also show that these cell preparations have low levels of *CD45*, a marker of white blood cells, compared to RNA isolated from opossum spleen (Supplementary Information Table 1).

## cAMP/MPA induce decidual regulator genes (40/40)

We assayed the response of MdESF to treatment with eutherian ESF differentiation media containing the cyclic AMP (cAMP) analogue 8-Br-cAMP and the progesterone (P4) analogue medroxyprogesterone acetate (MPA) (Fig. 1a), hereafter referred to as decidualizing stimuli. RNAseq of both stimulated and unstimulated MdESF revealed endogenous expression of numerous core regulatory genes critical to eutherian decidualization (Fig. 1b). From a curated list of 28 transcription factor (TF) genes with documented roles in decidualization (Extended Data Table 1), 22 are expressed in stimulated MdESF and 13 are significantly upregulated (p <0.05) (Fig. 1b and Fig. 2e). Seven decidualization TF genes are downregulated, though still expressed, and two TFs are unchanged in expression. Most notable, the upregulated gene set contains numerous TFs with well characterized roles in decidualization: *FOXO1*^*17,18*^*, PGR*^*19*^*, CEBPB*^*17,20*^*,HOXA10*^*21,22*^*, HOXA11*^*23,24*^*, GATA2*^*25*^*, ZBTB16*^*26,27*^*, KLF9*^*28-30*^*, HAND2*^*31*^*, STAT3*^*32,33*^, and *MEIS1*^*34*^ (Fig. 1b; Fig. 2e). In contrast to this conserved transcriptional regulatory response, classical markers of decidualization, *e.g. PRL*, *IGFBP1*, *CGA*, and *SST*, are neither expressed in unstimulated MdESF nor induced in response to decidualizing stimuli (Fig. 1c). We conclude that a substantial part of the DSC core GRN is also in place in opossum ESF and is responsive to progesterone and cAMP, but does not control a decidual phenotype.

**Figure 1.**
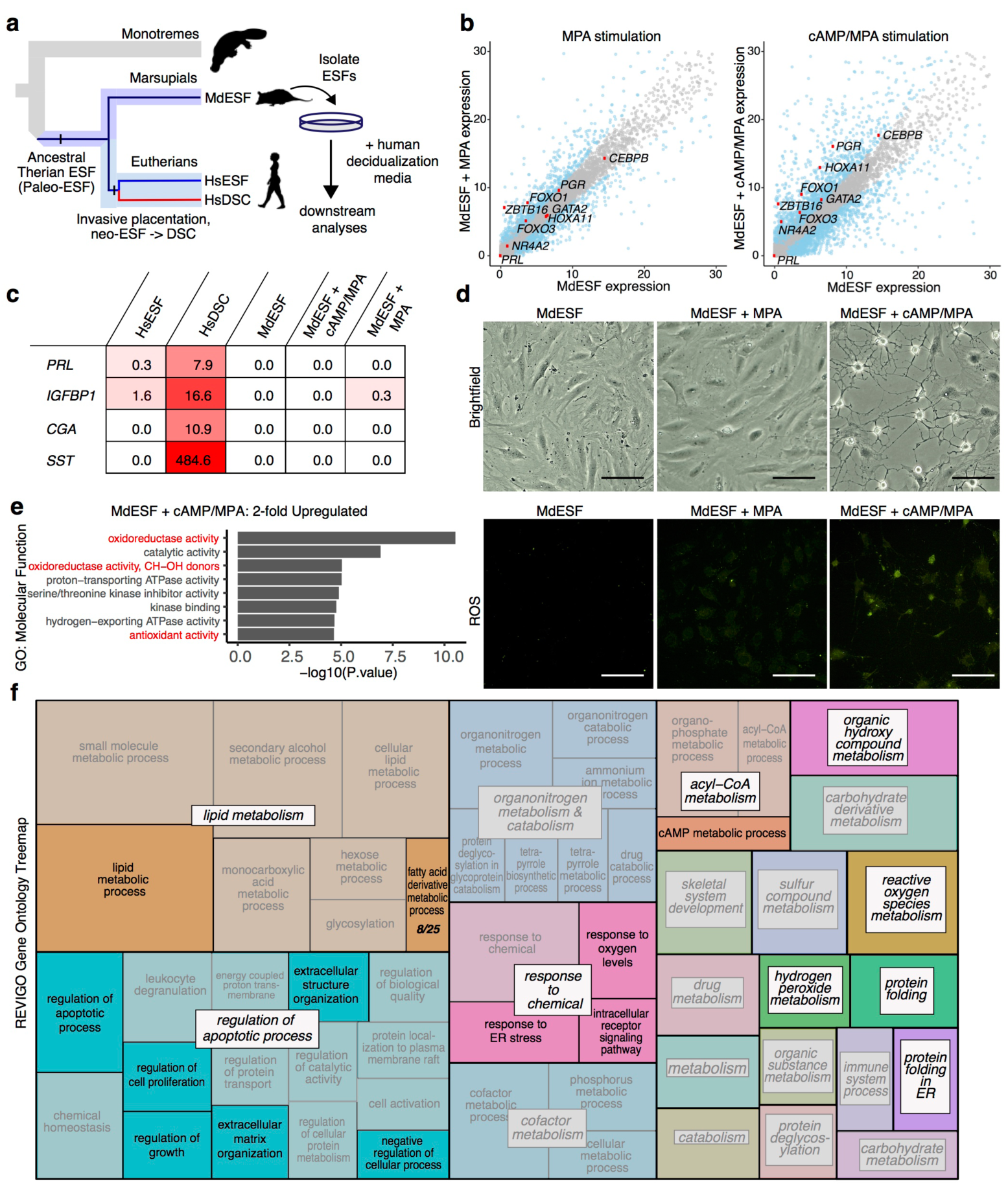
Transcriptomic and morphological response of MdESF to eutherian decidualizing stimuli. **a,** Experimental strategy and cell type phylogeny of mammalian ESF celltypes. ESF in crown group marsupials and placentals are homologous and descend from a common ancestral therian ESF. In eutherians, DSC arose from ancestral eutherian ESF and thus are sister cell type to eutherian ESF. **b,** Transcriptional response of core regulatory genes involved in eutherian decidualization in MdESF treated with either MPA or decidualizing stimuli. Blue dots represent significant differential expression relative to unstimulated MdESF (n=3, p-value<10^−6^). Grey dots represent no significant chance in expression. Each point represents the mean of three replicates. **c,** Gene expression heatmap (mean of three replicates in TPM) of classical decidual markers in HsESF, HsDSC, and MdESF treated with decidualizing stimuli. Mean of three replicates is shown. **d,** Morphological response of and ROS detection in unstimulated MdESF and MdESF stimulated with either MPA or decidualizing stimuli. ROS detection by treatment with H2DCFDA, a general oxidative stress indicator. Scale bars are 10uM. **e,** Differential expression of upregulated GO categories in MdESF treated with decidualizing stimuli relative to unstimulated control (n=3, adjusted *P* values for each GO term are shown). **f,** Visualization of GO term clusters associated with cAMP/MPA treatment in MdESF. Colored boxes represent semantic similiarity. Size of boxes represents p-value assigned to that cluster.

**Figure 2.**
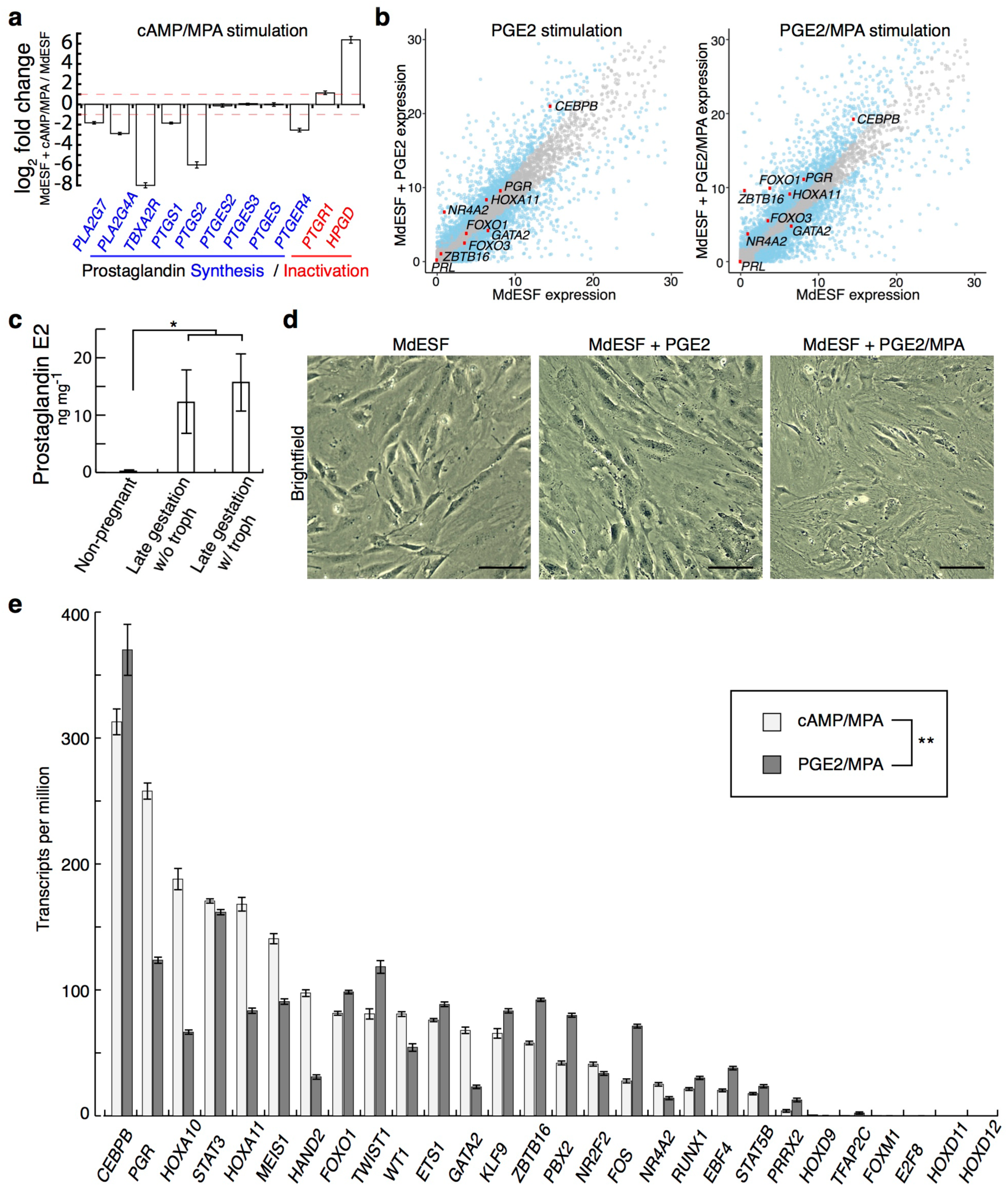
Transcriptomic and morphological response of MdESF to MPA and PGE2. **a,** Transcriptional response of KEGG pathway genes associated with prostaglandin signaling in MdESF treated with decidualizing stimuli relative to unstimulated control (n=3, log2 fold change shown). Red dashed line represents 2-fold change. **b,** Transcriptional response of core regulatory genes involved in eutherian decidualization in MdESF treated with either PGE2 alone or PGE2/MPA. Blue dots represent significant differential expression relative to unstimulated MdESF (n=3, p-value<10^−6^). Grey dots represent no significant chance in expression. Each point represents the mean of three replicates. **c,** Increased prostaglandin E2 (in ng per mg total protein) in pregnant (late gestation) versus non-pregnant *M. domestica* females (n=2 females per sample) as measure by ELISA (*, one-tailed t-test on log transformed data, p=0.014). **d,** Morphological response of unstimulated MdESF versus treatment with PGE2 or PGE2/MPA. Scale bars are 10 μM. **e,** Expression of 28 decidualization transcription factor genes in MdESF in response to treatment with either cAMP/MPA (black bars) or PGE2/MPA (grey bars). Non-parametric correlation of expression between treatments and across genes was found to be 0.863 (**, Spearman’s rho, p=3.46 10^−9^). Error bars are standard error of the mean.

## cAMP/MPA induce cell stress in MdESF (37/40)

Gene ontology (GO) enrichment analysis of differentially expressed genes after cAMP/MPA treatment revealed upregulation of genes associated with oxidative stress, mitochondrial stress, and apoptosis, as well as downregulation of genes associated with mitosis, DNA replication, and cytoskeletal organization (Fig. 1e and Extended Data Fig. 1a). Outwardly, stimulated MdESF exhibited a rapid morphological response suggestive of cytoplasmic architectural remodeling (Fig. 1d and Supplemental Video 1). The extent of this morphological response was dependent on both cAMP concentration and duration of treatment (Extended Data Fig. 1c). Remarkably, this morphological effect was reversible insofar as the cells reverted back to their normal morphology within 16 hours after withdrawal of decidualizing stimuli (Extended Data Fig. 1d). Gene ontology treemaps, which represent the function of genes and their differential expression in response to cAMP/MPA, supported the hypothesis that stimulated MdESF undergo a cellular stress response, as GO terms associated with endoplasmic reticulum (ER) stress, apoptosis, reactive oxygen species (ROS) metabolism, and protein folding response were significantly upregulated (Fig. 1f and Supplementary Information Fig. 4). In line with this observation, stimulated MdESF exhibited elevated levels of intracellular ROS relative to unstimulated cells or cells stimulated with MPA alone (Fig. 1d and Extended Data Fig. 1b). These data indicate that treating MdESF with decidualizing stimuli results in a rapid morphological response that is associated with increased intracellular ROS and the induction of genes counteracting oxidative stress, suggesting that, rather than leading to decidual differentiation, MdESF exposed to decidualizing stimuli undergo a classical stress response.

## PGE2 is likely a natural ligand of ESF (39/40)

Next we considered whether stress induced by treatment with decidualizing stimuli could be an artifact of treating cells with extracellular cAMP, rather than a natural ligand activating intracellular cAMP signaling. To address this, we sought a physiologically relevant signal that increases intracellular cAMP in these cells. Prostaglandin E2 signaling is of particular interest given that (1) PGE2 is able to induce decidualization via cAMP signaling in human and rodent ESF^11,13^, (2) the PGE2 receptor PTGER4 is widely expressed in ESF in mammals^16^, and (3) the recent finding that prostaglandin synthase (PTGS, a.k.a. COX2) and prostaglandin E synthase (PTGES) are both expressed in the opossum uterus after embryo attachment^35^. Furthermore, PGE2 is likely a key component of the inflammatory signaling from which the eutherian implantation reaction is derived^5,35,36^. In our cAMP/MPA stimulated cells we see a particularly striking effect on lipid metabolism, a critical pathway in the production of phospholipid-derived prostaglandins (Fig. 1f). Indeed, prostaglandin KEGG pathway genes were enriched in lipid metabolism, e.g. 33% of genes listed in fatty-acid derivative metabolic process are involved in prostaglandin metabolism (Fig. 1f). Furthermore, transcriptomic analyses of stimulated MdESF suggested that this treatment negatively regulates the predominant PGE2 receptor, *PTGER4*, as well as genes for synthesis of prostaglandins, *e,g, PTGS2* and *PTGES* (Fig. 2a), and positively regulates catabolic enzymes that function to degrade prostaglandins, *e.g. HPGD* and *PTGR1* (Fig. 2a). These data suggest stimulated MdESF “try to compensate” for the effect of cAMP by modulating the prostaglandin synthesis pathway.

A survey of RNAs present in unstimulated MdESF showed that only two prostaglandin signaling receptors, *PTGER4* and *TBXA2R,* are expressed in these cells (Extended Data Fig. 2d). In order to test whether PGE2 could be the natural ligand inducing intracellular cAMP signaling in these cells we assayed by RNAseq the response of MdESF to PGE2 with and without MPA. KEGG pathway genes involved in prostaglandin synthesis and inactivation exhibited similar differential regulation in response to PGE2/MPA as do decidualizing stimuli, suggesting similar regulatory responses by MdESF (Extended Data Fig. 2b,c). Uterine tissue from pregnant and non-pregnant *M. domestica* showed substantially higher amounts of PGE2 in pregnant females versus non-pregnant females, suggesting that PGE2 increases *in utero* during gestation (Fig. 2c). Contrary to treatment with decidualizing stimuli, MdESF treated with PGE2 and with or without MPA did not exhibit a readily apparent dendritic phenotype (Fig. 2d). Nevertheless, PGE2/MPA-treated MdESF do show elevated levels of intracellular ROS (Extended Data Fig. 2e).

We next investigated the effect of PGE2/MPA on the expression of core decidualization regulatory genes (Fig. 2b,e). Remarkably, all 22 decidual TF regulatory genes expressed in cAMP/MPA-stimulated MdESF cells are also expressed in PGE2/MPA-stimulated cells (Fig. 2e), showing a marked non-parametric correlation in expression levels (Spearman’s rho=0.863, p=3.46 10^−9^), and 12 of the 13 upregulated genes under cAMP/MPA are also upregulated with PGE2/MPA. We conclude that responding to PGE2 signaling is a physiological part of MdESF biology and that PGE2/MPA also regulates the expression of the same TF network as cAMP/MPA treatment. Moreover, the results suggest that this PGE2-induced TF network is homologous to that activated during the differentiation of human DSC.

## PGE2-induced stress response in MdESF (38/40)

We next asked if, as seen in cells treated with decidualizing stimuli, PGE2/MPA treatment also induced a stress response in MdESF. We surveyed differential expression of genes involved in oxidative stress response, apoptosis, and ROS-associated endoplasmic reticulum stress (unfolded protein response, or UPR) (Extended Data Fig. 3). MdESF treated with both cAMP/MPA and PGE2/MPA significantly upregulated genes associated with counteracting oxidative stress, including *GCLM*, *GPX3*, *GPX4*, *SOD1, SOD3,* and *CAT* (Extended Data Fig. 3a). Both treatments upregulated the apoptotic genes *BCL2L11 (BIM)* and *GADD45A* (Extended Data Fig. 3b). In contrast to treatment with PGE2/MPA, cAMP/MPA induced a distinct stress response in genes associated with UPR and TRAIL-related apoptosis, upregulating *ERN1 (IRE1), HSPA5, HSP90B1, HSP90AA1, CALR,* and *TNFSF10* (Extended Data Fig. 3b,c). Lastly, to determine if clusters of genes associated with oxidative stress and apoptosis were differentially expressed in both cAMP/MPA and PGE/MPA, we analyzed GO term clusters shared between the treatments. This analysis also suggested a shared stress response in MdESF treated with decidualizing stimuli or PGE2/MPA, in which GO terms associated with stress and inflammation, e.g. “regulation of reactive oxygen species metabolism”, “protein folding”, and “leukocyte degranulation”, were shared between these treatments (Fig. 1e and Supplementary Information Fig. 7). Similarly, PGE2-alone and PGE2/MPA shared GO terms specifically related to “hypoxia”, “autophagy”, and “regulation of cell death” (Supplementary Information Fig. 6 and Supplementary Information Fig. 7). These results suggest that PGE2/MPA, signals that are present in the pregnant opossum uterus, induce a stress response similar to treatment with cAMP/MPA, including elevated levels of intracellular ROS (one-tailed t-test on log transformed data, p=0.0186), but without the dendritic morphological response (Extended Data Fig. 2e). We conclude that the *in vitro* cAMP/MPA-induced stress reaction is mimicked with the more physiological PGE2/MPA treatment. Further, these data are consistent with a conserved role for PGE2 during pregnancy of therian mammals.

## Subcellular localization of FOXO1 (34/40)

In many cells FOXO TFs generically function in stress response, counteracting oxidative stress and apoptosis, as well as regulating gluconeogenesis and glycolysis^37^. FOXO1 also is an early acting TF in the differentiation of human DSC^38^. Therefore, we sought to compare how *FOXO1* mRNA and FOXO1 protein stability and sub-cellular localization are regulated in opossum ESF. As observed in human ESFs, *FOXO1* RNA is present in unstimulated MdESF, but FOXO1 protein is absent^16,39^, likely due to Akt-dependent proteasomic degradation^37^ (Fig. 3a). In response to MPA treatment, FOXO1 accumulates in the cytoplasm (Fig. 3b), suggesting MPA alone can counteract the proteasomic FOXO1 degradation. Decidualizing stimuli and treatment with PGE2/MPA resulted in nuclear translocation of FOXO1 as well as cytoplasmic loading (Fig. 3c,d,e), suggesting cAMP/PKA signaling controls FOXO1 nuclear localization. In response to induction of oxidative stress, FOXO1 behaved similarly to cAMP or PGE2 treatment, suggesting that post-translational modifications of FOXO1 act as a sensor of oxidative stress in MdESF (Fig. 3f). Moreover, immunofluorescence on uterine sections from pregnant *M. domestica* females in the late stages of gestation (11.5 dpc) found nuclear FOXO1 in uterine stromal cells near the luminal epithelium, suggesting that FOXO1 activation is part of the physiological role of MdESF during pregnancy (Fig. 3h). Strikingly similar results were obtained for FOXO1 in human DSC (Extended Data Fig. 4a), suggesting post-translational regulatory control of FOXO1 as found in human DSC in response to decidualizing stimuli has been inherited from the ancestral paleo-ESF and ancestrally was part of a PGE2-induced cell stress response.

**Figure 3.**
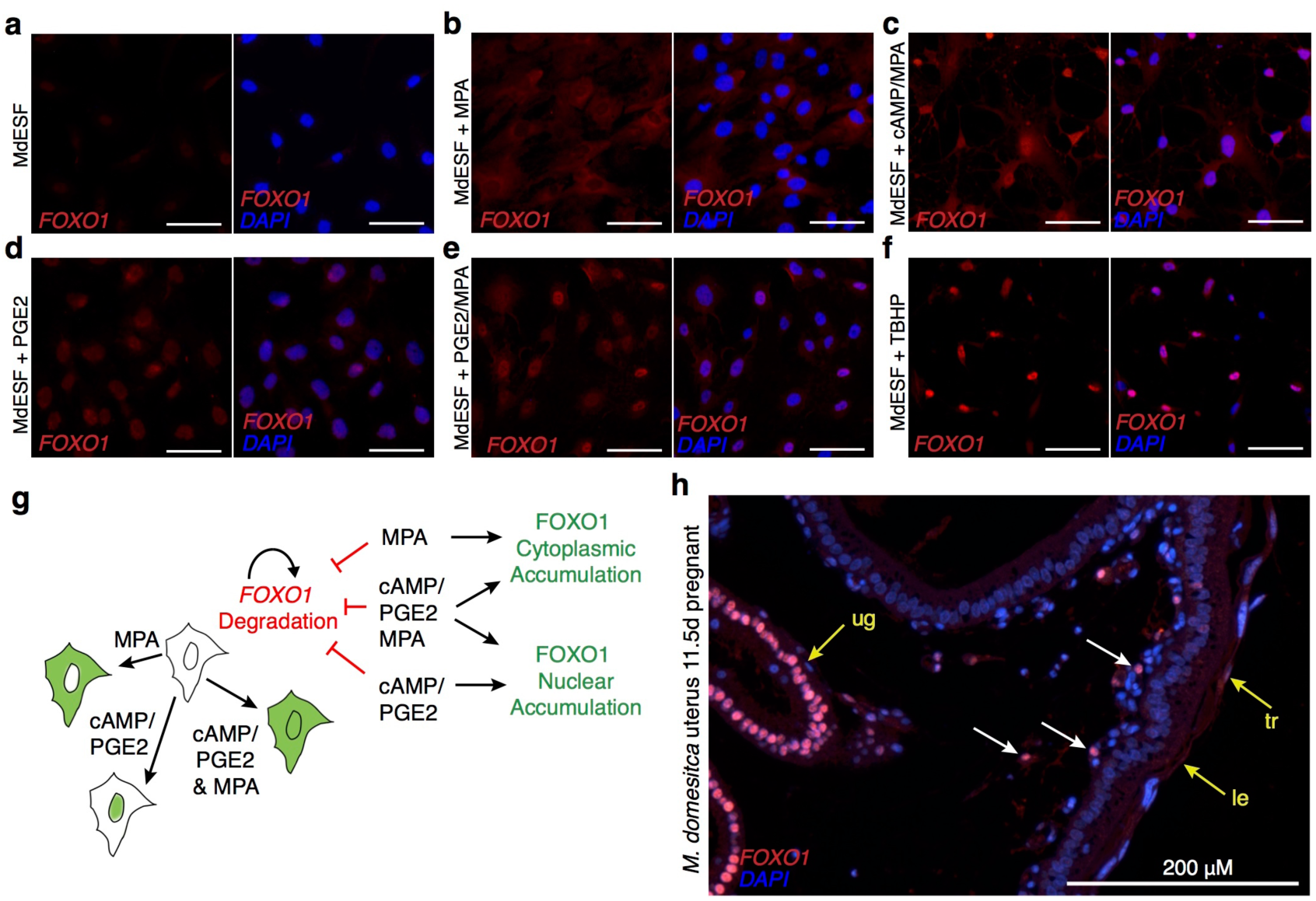
Post-translational regulatory control of FOXO1 by PGE2 and oxidative stress in MdESF. **a-h,** Immunofluorescence of FOXO1 in MdESF and pregnant *M. dometica* uterus. **a,** FOXO1 is not detected above background in unstimulated MdESF. **b,** FOXO1 protein is detected in the cytoplasm, but not in the nucleus, in MdESF treated with MPA alone. **c,** FOXO1 translocates to the nucleus and is detected in the cytoplasm in MdESF treated with decidualzing stimuli cAMP/MPA. **d, e,** FOXO1 is detected in the nucleus and cytoplasm in MdESF treated with either PGE2 alone or PGE2/MPA. **f,** Oxidative stress induces FOXO1 to translocate to the nucleus in MdESF treated with tert-butyl hydrogen peroxide (TBHP). Scale bars are 10 M. **g,** Model showing post-translational regulatory control of FOXO1 in MdESF treated with decidualizing stimuli or cAMP, MPA, PGE2 alone. **h,** FOXO1 immunofluorescence of uterine tissue in cross section during late gestation (11.5 d.p.c.). All samples counterstained with DAPI. Arrows indicate FOXO1 detection. ug, uterine glands; tr, trophoblast; le, luminal epithelium.

## FOXO counteracts apoptosis in opossum ESF (40/40)

In order to assess the functional role of FOXO1 activation in opossum ESF we assayed oxidative stress and apoptosis by treatment with H2DCFDA, to detect ROS, and propidium iodide (PI), to detect early stages of apoptosis, in stimulated and unstimulated MdESF (Fig. 4). Apoptosis, i.e. PI staining, was markedly elevated in MdESF treated with decidualizing stimuli (Fig. 4a,b; 4 fold increase, one-tailed t-test p=2.0 10^-5^). That was also the case for PGE2/MPA but to a lesser degree (1.8 fold increase, one-tailed t-test p=0.015). We transfected MdESF with siRNAs targeting *FOXO1* and *FOXO3* RNA transcripts. We confirmed depletion of *FOXO1* RNA (as well as *FOXO3* RNA) by qPCR and depletion of FOXO1 protein by Western blot (Extended Data Fig. 4b,c). siRNA mediated KD of *FOXO1* and *FOXO3* increased signals for apoptosis (ANOVA on log transformed fluorescence values, KD effects FOXO1=2.4 fold, FOXO3=1.8 fold, overall ANOVA p=5.91 10^-8^), suggesting they function additively in protecting against apoptosis, *i.e.* there was no statistical interaction effect (Fig. 4a,b). However, there is no significant effect of FOXO KD on ROS levels in our data, suggesting that ROS production in response to decidualizing stimuli is not regulated by FOXO proteins.

**Figure 4.**
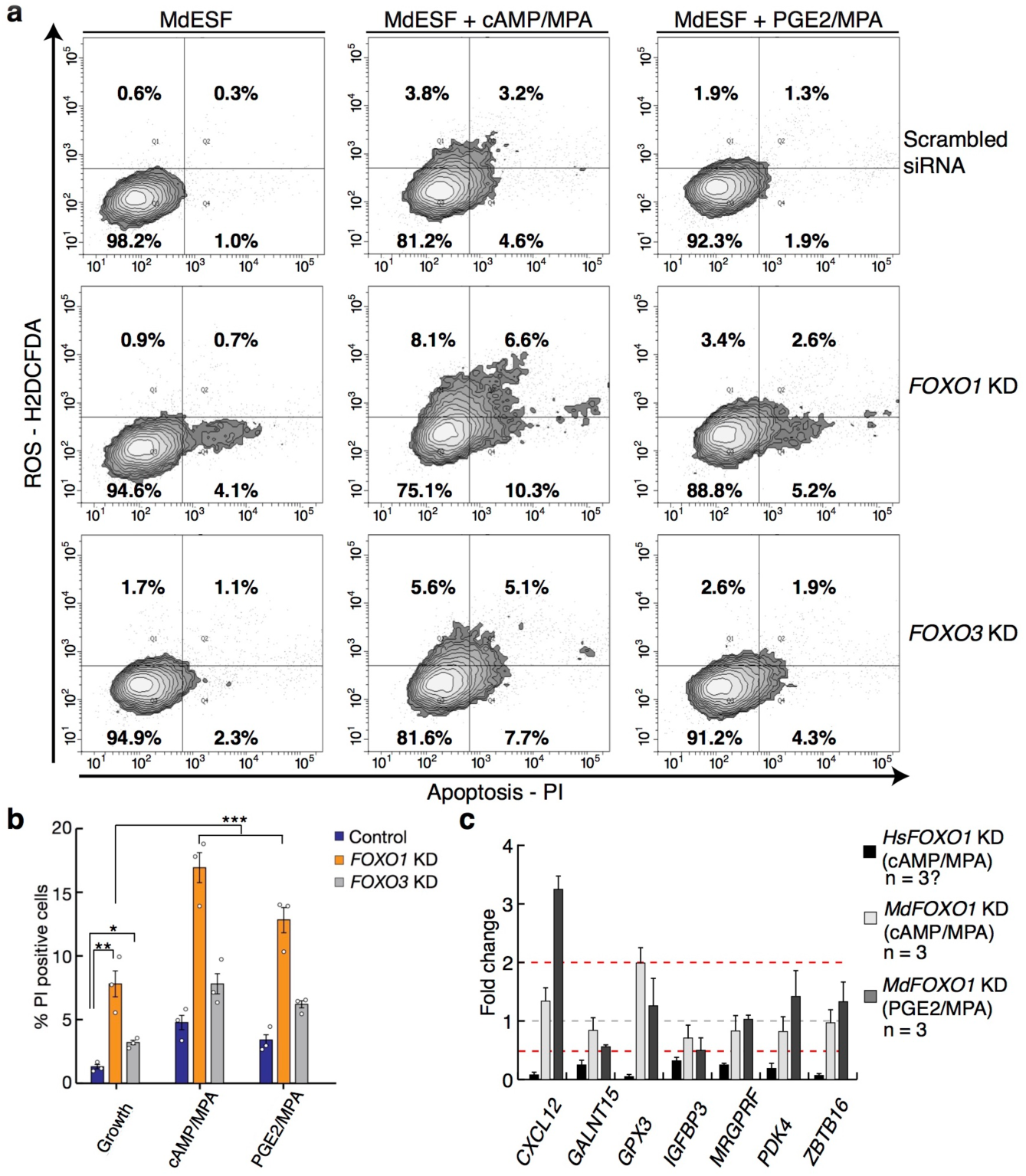
FOXO proteins counteract apoptosis in stimulated MdESF. **a,** Treatment of MdESF with cAMP/MPA or PGE2/MPA results in more cells positive for ROS and apoptosis, PI staining (top panels). Knockdown of *FOXO1* in unstimulated MdESF increases cell counts for apoptotic cells relative to control treated with scrambled siRNA control (middle, left panel). Knockdown of *FOXO1* and treatment with cAMP/MPA or PGE2/MPA significantly increased cell counts for apoptosis along with a moderate increase in ROS (middle panels). Knockdown of *FOXO3* in unstimulated MdESF increases cell counts for ROS and apoptosis (bottom, left panel). Knockdown of *FOXO3* and subsequent treatment with decidualizing stimuli or PGE2/MPA results in increased cell counts for ROS and apoptosis (bottom panels). Quadrants are set to maximum extent of boundaries in unstimulated MdESF treated with scrambled siRNA. Q1, ROS positive only; Q2, ROS and PI positive; Q3, negative for ROS and PI; Q4, negative for ROS and positive for PI. Percentage in each quadrant represents mean of three replicates. **b,** Bar graph showing mean percent positive cells for quadrants 2 (ROS and PI positive) and 4 (PI positive only) (n=3, 2-tailed t-test, *, p<0.05, **, p<0.005; ANOVA on log transformed fraction of cells, ***, p=5.91 10^−8^). Blue, scrambled siRNA control; orange, *FOXO1* knockdown; gray, *FOXO3* knockdown. Empty circles represent individual data points for each replicate.**c,** Fold change of *FOXO1*-responsive genes in human DSC in MdESF stimulated with either cAMP/MPA or PGE2/MPA in *FOXO1* perturbation background relative to stimulated control treated with scrambled siRNA. Data are the average of three replicates  standard error of the mean.

*FOXO1* knockdown resulted in marginally more cells positive for apoptosis than did *FOXO3* perturbation (Fig. 4b; 1.31 fold, p=0.045). These results suggest that the ancestral function of FOXO genes in paleo-ESF was similar to the highly conserved, pan-metazoan roles of FOXO genes in classical stress response^40^. Therefore, we hypothesized that the regulatory linkages downstream of FOXO1 have diverged since the eutherian-metatherian split. From a list of genes in human DSC that are positively regulated by FOXO1 (*i.e.* decrease in expression after FOXO1 KD), we randomly selected seven genes that are also upregulated in MdESF stimulated with cAMP/MPA and PGE2/MPA. Of the seven genes one, *IGFBP3*, significantly decreased in expression and all other genes either did not respond or increased in expression in response to *FOXO1* KD in MdESF (Fig. 4c). This result suggests that FOXO1 transcriptionally regulates distinct sets of genes in MdESF and HsDSC. If we assume that the reproductive mode in MdESF is characteristic of the ancestral paleo-ESF, these data also suggest that the evolution of mammalian DSC proceeded through modifying the target gene set of a largely conserved core GRN that includes FOXO1.

## Discussion

Here we show that, whereas human ESF respond to decidualizing stimuli with a compensated physiological phenotype, the DSC, opossum ESF exhibit a classic stress phenotype. This difference was also found, though to a lesser degree, when we used a more physiological signal, prostaglandin E2 (PGE2) instead of extracellular cAMP. The response of human and opossum ESFs was remarkably similar at the level of DSC regulatory gene expression, both in terms of transcriptional as well as post-translational regulation as in the case of FOXO1. While post-translational activation of FOXO1 is necessary in human cells for the expression of DSC effector genes, e.g. *PRL*, *IGFBP1*, etc., in opossum ESF the functional role of activated FOXO1 and FOXO3 is to protect the cells against apoptosis.

We propose that the signaling pathway and large parts of the TF network are homologous and to some degree conserved between eutherian DSC and marsupial MdESF, suggesting that these components were also present in paleo-ESF. Further, we hypothesize that there were at least two distinct molecular changes that led to the evolution of DSC. On one hand we find a small number of decidual TFs that are not expressed in stimulated MdESF, *viz*. members of the 5’ HoxD cluster *HoxD12*, *HoxD11* and *HoxD9*, as well as *FOXM1*, *TFAP2C*, *PRRX2* and *E2F8*. Thus, *cis-*regulatory linkages of core decidual regulators likely accumulated in response to increased intracellular cAMP to accommodate the novel functions of DSC. For example, in this list *E2F8* functions in regulating a polyploidization^41^, which is a derived feature of DSC. On the other hand, we also find that DSC effector genes of the conserved TF network are different. Thus, another element by which the ancestral ESF cell type evolved into DSC was the “rewiring” of gene regulation downstream of FOXO1 and other decidual regulatory genes, *e.g. CEBPB*, *PGR*, *HOXA10*, *HOXA11*, and *GATA2* (Fig. 5). We tested this model by comparing the effect of FOXO1 KD on effector gene expression and found that in fact the regulatory role of human FOXO1 in these cells is extensively different from that in opossum ESF. Exactly how this “downstream reprogramming” was effected in evolution is not known and needs to be the subject of further investigation.

**Figure 5.**
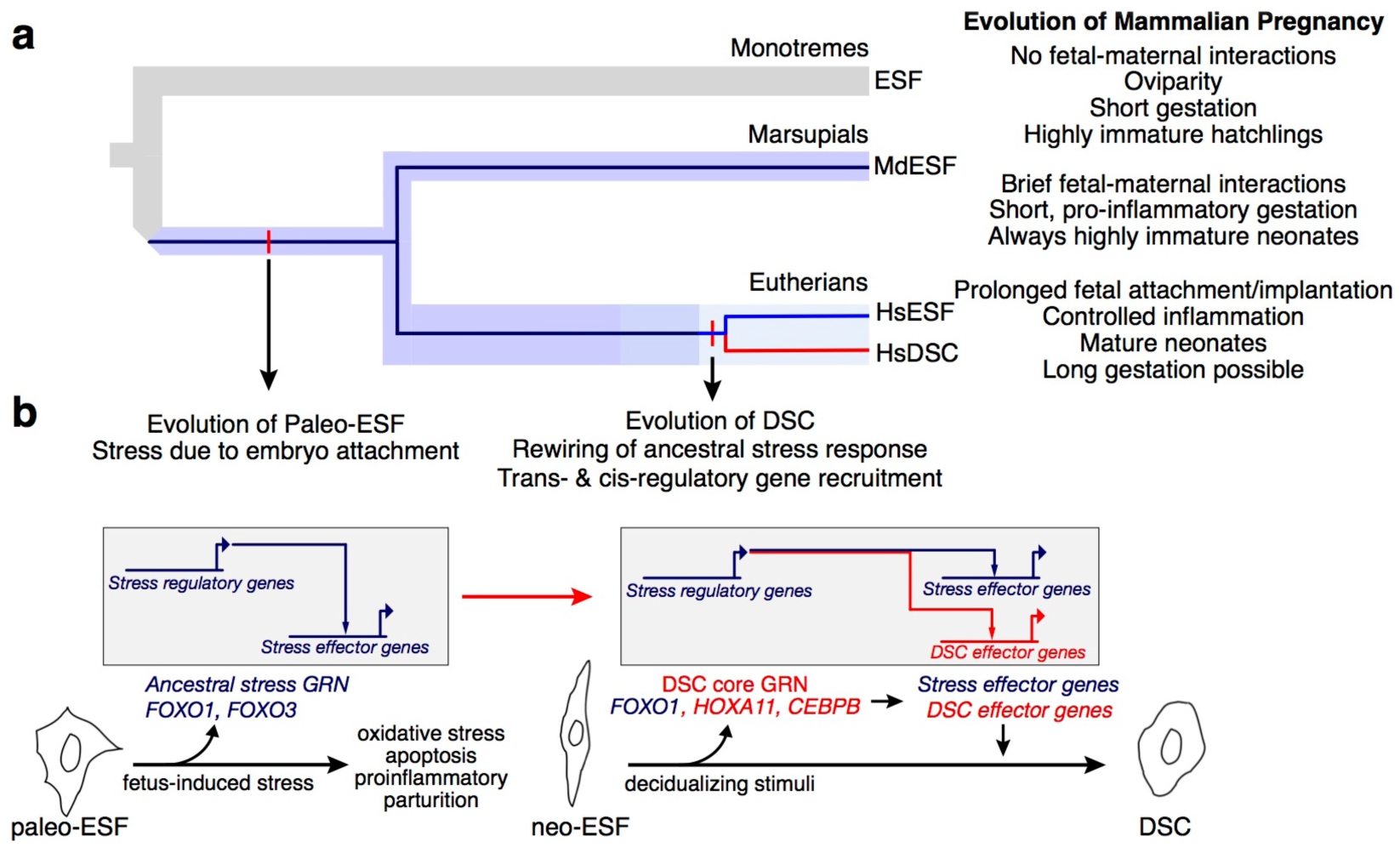
The eutherian mammal decidual cell type evolved from a cellular stress response. **a,** Phylogeny showing the three major mammalian clades. Embedded in the species tree, the cell type tree for ESF and DSC cell types is drawn. Paleo-ESF is an ancestral cell type to crown group neo-ESF, such as human HsESF, which arose after the split of metatherian (marsupials) and eutherian (placental) mammals. The eutherian neo-ESF is defined by its ability to differentiate into DSC, which is the sister cell type to eutherian neo-ESF. **b,** Proposed model of the evolution of eutherian DSC from a paleo-ESF ancestral cell type. Paleo-ESF underwent a stress response during the short gestational period in which fetal-maternal interactions lead the proinflammatory, stress-induced parturition seen in crown marsupials. The ancestral stem eutherian rewired the stress-related regulatory modules to stabilize an alternative gene regulatory state and control the expression of effector genes, which allow fetal implantation and invasive placentation, long gestation, and anti-inflammatory phenotypes typical of crown eutherian mammals.

An alternative explanation for our results is that the decidual cell type may have evolved in the stem therian lineage, i.e. before the most recent common ancestor of eutherians and marsupials, but has been lost in marsupials. Marsupial reproduction shares many plesiomorphic reproductive traits with the most basal branching mammals, the monotremes^35^. Although opossums do not lay eggs, along with the platypus they have a relatively short gestation period and give birth to highly altricial young^42,43^. Further, DSCs are necessary for maintaining pregnancy in species with the kind of invasive placentation that is only found in eutherian mammals^8^. For these reasons, it is likely that the DSC is truly a specific trait of eutherian mammals and not one with an older origin that was subsequently lost in the marsupial group.

Our model of stress-derived decidual differentiation explains a number of otherwise puzzling facts. First, there is evidence that during physiological decidualization in humans the stromal cells produce intracellular ROS that mediate the decidualization process ^44-47^. Here we show that genes responsive to decidualization stimuli in a distantly-related ESF cell type are also intimately linked to stress-related decidualization in eutherians^44^, e.g. *GPX3, SOD1, SOD3,* and *CAT,* and more interestingly these genes are also associated with a stress coping mechanism. Further, we show that apoptosis-related genes known to play a role in decidualization, e.g. *GADD45A, TRAIL,* and *BCL2L11*^44,48^, also are upregulated in MdESF. In the context of these stress-related genes, we also show that FOXO1 and FOXO3 act as an oxidative stress sentinel that counteracts apoptosis. Moreover, decidualization is associated with endoplasmic reticulum stress ^49^, which we also observe in genes specifically associated with ER stress and UPR in opossum cells treated only with cAMP/MPA, suggesting that a derived aspect of eutherian ESF is to cope with increased levels of intracellular cAMP. Finally, a sub-population of endometrial stromal cells undergoes cellular senescence in humans, a senescent phenotype that plays a critical role in implantation^50^. Thus, peculiar features of human decidualization, *e.g.* redox signaling, ER stress, cell senescence, are readily explained by our model, wherein decidual differentiation mechanisms arose in evolution from a pregnancy-related stress response that consequently activates many of the same regulatory genes and physiological processes as activated in decidual cells during differentiation.

In evolution, structural and developmental changes can result in cells being exposed to drastically altered or novel environments. In mammals, the evolution of pregnancy and in particular the evolution of extended gestation resulted in the endometrium functioning under exposure to a range of new stimuli, and with new requirements for the success of reproduction. As these developmental changes occurred, it is reasonable to expect that exposure of cell types to novel tissue and developmental environments can be a source of cellular stress. These anciently conserved pathways can be a rich source of hardwired, modular components of stress GRNs. In this case, evolution can mitigate cellular stress in a variety of ways, including decreasing the stress-inducing stimulus, but also through co-option of the stress pathways into normal physiological function. Our data suggest that the decidual stromal cell type has evolved through genetic assimilation of a stress response that is likely directly related to the invasion of trophoblast into maternal tissues, the condition seen in crown eutherians today.

While the evidence presented here pertains only to the special case of mammalian decidual cells, the recent discovery that ROS play the role of physiological signaling molecules during the differentiation of other cell types, e.g. neurons^51,52^ and mesenchymal cells^53^ (and functioning in tumorigenesis), may indicate that the derivation of novel cell types from stress responses might be a more general phenomenon. The evolution of novel cell types, with novel physiological functions, could therefore be understood as fulfilling physiological needs for which the ancestral body plan or developmental program cannot compensate. The evolutionary rewiring of stress responses could be a broadly applicable model for cell type origination in evolution.

## Methods

### Animal husbandry

All animal procedures were conducted under protocols approved by the Institutional Animal Care and Use Committee (#11313) of Yale University. Opossum uterine tissue was collected from a *M. domestica* colony housed at Yale University. For ESF isolation, uterine tissue was harvested from a non-pregnant *M. domestica* female. For immunohistochemistry and ELISA, tissue from specific stages of the reproductive cycle were collected by following a standard breeding protocol outlined in Kin et al.^16^. Once collected tissue was stored for immunohistochemistry analysis in 4% paraformaldehyde in PBS 24 h, then washed in 50% ethanol 1 h, then twice in 70% ethanol 1 h then stored in 70% ethanol at -20 °C. Tissue was stored for western blot and ELISA analysis by snap freezing in liquid nitrogen and then stored at -80°C.

### ESF isolation from the uterus of Monodelphis domestica, cell line validation, and propagation

MdESF were isolated as previously described^16^. Briefly, primary endometrial stromal fibroblasts were harvested by enzymatic digestion and centrifugation combined with Percoll density gradient. The uterus of an adult female grey short-tailed opossum *M. domestica* was dissected, cut in half longitudinally, and cut into 2-3 mm fragments. These were digested with 0.25% trypsin-EDTA for 35 min at 37°C and digested in Dissociation Buffer (1 mg/ml collagenase, 1 mg/ml Dispase, 400 μg/ml DNaseI) for 45 minutes at room temperature. Cell clumps were subsequently homogenized by passage through a 22-gauge syringe. Passage through a 40 μm nylon mesh filter removed remaining fragments. This lysate was used to generate a density gradient by centrifugation at 20,000 g for 30 min. A single cell suspension was generated from this lysate and was subsequently layered onto a Percoll gradient (1.09g/cc Percoll, GE Healthcare Life Sciences) and centrifuged at 400 g for 20 min to allow for cells to settle to their respective density layers. Using a 25-gauge needle, each 1 ml layer was removed working from low to high density and washed into 5 ml 50 mM NaCl. As with previous iterations of this protocol in this laboratory, layers 6 thru 8 generally contained a fairly homogeneous population of cells that outwardly exhibited fibroblast characteristics. Cells in these layers were pelleted, resuspended in growth media, and cultured in 24 well plates. To facilitate enrichment of fibroblasts versus epithelial cells, media was exchanged in each well after two hours in order to remove floating cells that had not yet attached. To validate our cell line, we conducted comparative qPCR and immunofluorescence for proteins that mark either epithelial or mesenchymal (fibroblast) cells. Transcription factors indicative of ESFs were enriched in RNA from Percoll layer 8 relative to later 3 or RNA isolated from spleen (Supplementary Information Table 1). Immunofluorescence on cells from Percoll layer 8 showed enrichment of the mesenchymal protein VIMENTIN and no epithelial contamination as judged by expression of CYTOKERATIN. (Supplementary Information Fig. 1). Therefore, cells from later 8 were used in these experiments and were propagated in a T75 culture flask by sub-passaging over a period of 2 months at 33°C. Once confluent, cells were sub-passaged using Accutase (AT104, Innovative Cell Technologies) or a cell scraper. The experiments detailed here were conducted with passages 12-20.

### Cell culture, RNAseq, and transcriptomic analyses

MdESF were cultured in growth media with no antibiotic-antimycotic, containing (per liter): 15.56 g DMEM/F-12 (D2906, Sigma Aldrich), 1.2 g sodium bicarbonate, 10 ml sodium pyruvate (11360, Thermo Fisher), 1 ml ITS (354350, VWR), 100 ml charcoal-stripped fetal bovine serum (100-119, Gemini). Media was replaced every 4 days unless otherwise stated. Over the duration of these experiments, MdESF were found to be mycoplasma free (Supplementary Information Fig. 8), as shown by periodic PCR assays for mycoplasma contamination (30-1012K, Universal Mycoplasma Detection Kit, ATCC). For decidualizing stimuli, MdESF were cultured in growth media supplemented with cAMP-analgoue 8-bromoadenosine 3’-5’-cyclic monophosphate sodium sale (B7880, Sigma Aldrich) and progesterone-analgoue medroxyprogesterone 17-acetate (MPA) (M1629, Sigma Aldrich), at final concentrations of 0.5 mM and 1 μM respectively. Growth media was supplemented with prostaglandin E2 (14010, Cayman Chemicals) at a final concentration of 10 μM. For RNA sequencing of unstimulated and stimulated MdESF, cells were grown to 70% confluency in T75 flasks and treated with the respective stimuli for two days prior to harvesting with Buffer RLT and subsequent processing with RNeasy Mini Kit (74104, Qiagen) following the manufacturer’s protocol. Illumina sequencing libraries were generated from RNA by Poly-A selection and sequenced by the Yale Center for Genome Analysis on the Illumina Hiseq 2500 system. For transcriptomic and gene ontology enrichment analyses, see below.

### Immunocytology

MdESF or HsESF were grown in a 8-well chamber slide (12-565-18, Fisher) to 70% confluency. After treatment, cells were fixed with 4% paraformaldehyde in PBS for 15 min at room temperature. Cells were washed 2x in ice-cold PBS, subsequently incubated for 10 min in 0.25% Triton X-100 in PBS, and finally washed 3x for 5 min/wash in PBS. A blocking solution was applied with 1% bovine serum albumin (BSA), 0.25% Triton X-100 in PBS for 30 min at room temperature. Cells were then incubated in blocking solution at 4°C overnight in the following primary antibodies: 1:200 rabbit anti-cytokeratin (ab9377, Abcam); 1:200 mouse anti-vimentin (sc-6260, Santa Cruz); mouse anti-FKHR (FOXO1) (sc-374427, Santa Cruz). Cells were subsequently washed the next day 3x for five minutes each in PBS and secondary antibody incubation was for one hour at room temperature in the dark. Secondary antibodies used in this study are: 1:200 Alexa Fluor 555 goat anti-mouse IgG (A21422, Thermo Fisher); 1:200 Alexa Fluor 488 goat anti-rabbit IgG (A11008, Thermo Fisher). Cells were then washed 3x for five minutes each in PBS and nuclei were stained with DAPI (10236276001, Roche). Finally cells were washed 1x for 5 min in PBS and observed with an Eclipse E600 microscope (Nikon) equipped with a Spot Insight camera.

### Western blot analysis

MdESF or HsESF were cultured in T75 flasks to 80% confluency. Cells were rounded up with Tryple and homogenized in RIPA buffer (89900, Thermo Fisher) supplemented with HALT Protease Inhibitor Cocktail (PI87785, Thermo Fisher) for 15 min. Suspensions were centrifuged for 15 min at 13,000 rpm at 4°C. Protein concentrations were determined with Pierce BCA Protein Assay Kit (23225, Thermo Fisher). Cell lysates were diluted to achieve a solution with 30-60 μg total protein, combined with an equal volume of 2x NuPage LDS Sample Buffer (NP007, Thermo Fisher) with 2x NuPage Sample Reducing Agent (NP0004, Thermo Fisher), heated to 70°C, loaded in a NuPage 4%-12% Bis-Tris gel (NP0321BOX, Thermo Fisher), and electrophoresed at 130 volts for 60-90 min. Proteins were transferred to polyvinylidene difluoride membranes with the iBlot Gel Transfer System (Thermo Fisher). Membranes were subsequently incubated for one hour at room temperature in blocking buffer (3% BSA in PBST) and incubated with primary antibodies, listed above, overnight at 4°C. Primary antibody dilutions were the following: 1:200 FOXO1, VIMENTIN, and CYTOKERATIN. After primary incubation, membranes were then washed 3x for five min each in PBST and subsequently incubated with the corresponding HRP-conjugated secondary antibody, 1:5000 of either goat anti-mouse (sc-2005, Santa Cruz) or goat anti-rabbit (sc-2054, Santa Cruz). Signal was detected by incubating membranes in Clarity Western ECL substrate (1705060, Bio-Rad) in the dark for 5 min and visualized with a Bio-Rad Gel Doc System. Uncropped Western blots are provided in Supplementary Information Fig. 2 and Fig. 3.

### Immunohistochemistry

Uterine tissue was dehydrated through a graded ethanol series, cleared in toluene, and then embedded in paraffin. We cut 7 μm cross sections on a microtome and mounted on Shandon polysine precleaned microscope slides (6776215Cs, Thermo Fisher). Sections were stored in the dark at room temperature until they were stained. We localized the expression of FOXO1 using immunohistochemistry. Slides were de-parafinized in three successive washes of xylene (3 min each), then three successive washes of ethanol (3 min). Antigen retrieval was performed in citrate buffer (12 mM sodium citrate, pH 6.0, 98 °C, 1 hour). Endogenous peroxidase activity was blocked with Dako Peroxidase Block (Dako, 30 min). Slides were then incubated in primary antibody overnight. We used a goat polyclonal antibody generated against the N-terminal of human FOXO1 protein (1:5000 dilution, FKHR antibody sc-9809, SantaCruz). On day two slides were incubated in a donkey anti-goat IgG-HRP secondary (1 hour, 1:200 dilution, sc-2056 SantaCruz). Slides were then rinsed in PBS (5 min), PBS-BSA(0.1 %, 5 min), then incubated in TSA Plus Cyanine 3 system (1:50, 5 m, NEL744001KT, PerkinElmer Inc.). Slides were again washed in PBS (5 min) and PBS-BSA (0.1%, 5 min), counter stained in DAPI (1x, 2 min, 10236276001; Roche), washed in H2O (deionized, 5 min), and mounted in glycerol (50%).

### Prostaglandin E2 ELISA

Snap frozen uterine tissue from adult female *M. domestica* was homogenized in extraction buffer (0.1 M phosphate, 1 mM EDTA, pH 7.4) containing indomethacin (10 μM final concentration) using a mechanical homogenizer (TissueRuptor, QIAGEN). Cellular debris was removed by centrifugation (> RPM, 4 °C, 10 min). Tissue lysates were aliquoted into single use tubes and frozen at -80 °C. Protein concentration was measured using Pierce BCA Protein Assay Kit (23225, Thermo Fisher). After determining protein concentration, 1 mg of protein for each sample was used in the first round of ELISA against prostaglandin E2 following the manufacturer’s protocol (514010, Cayman Chemical). Each sample was run in duplicate at two different dilutions. A dilution series of PGE2 standard provided by the manufacturer was included in each run, and PGE2 values in ng per mg protein were calculated from these standards. Due to the sensitivity of this assay, some samples contained excess PGE2 and therefore required additional dilutions in order to fall within the calculable range of the standard. For these samples with excess PGE2, an additional ELISA plate was run with two additional dilutions. Samples were incubated in the provided PGE2 monoclonal antibody ELISA plate overnight at 4°C. The following day, the wells were washed 3x and the staining reaction was allowed to proceed for 45 minutes with shaking at 400 rpm on an orbital shaker in the dark. The plate was read at 405 nm on a Viktor X multilight plate reader (Perkin Elmer).

### Quantitative PCR

MdESF were cultured in T25 culture flasks to 70% confluency and subsequently transfected with siRNAs as above. After two days, cells were treated either with cAMP/MPA or with PGE2/MPA for four days. The change of media was accompanied by an additional round of siRNA transfection per above. At the time of RNA harvest, media was removed, cells were washed 2x in PBS, and cells were lysed directly in the flask with Buffer RLT Plus + beta-mercaptoethanol. RNA was harvested according to the manufacturer’s protocol (74034, RNeasy Plus Micro Kit, Qiagen). Reverse transcription of 3 μg of RNA was carried out with iScript cDNA Synthesis Kit (1708891, Bio-Rad) with an extended transcription step of three hours at 42°C. For qPCR, all reactions were carried out with Power SYBR Green PCR Master Mix (4368708, Thermo Fisher) in triplicate with 40 ng of cDNA for template in each technical replicate reaction. Fold change was calculated by finding the ddCt values relative to the expression of TATA Binding Protein. All qPCR primer sets were validated by analysis of melting curves for 2 different sets of primers for the same gene. Primer sets used in this study are listed in Supplemental Information Table 2.

### RNA-interference

MdESF at 70% confluency in 6 well culture plates were transfected with 25 nmol final concentration siRNAs targeting *FOXO1* (Mission custom siRNA, V30002, Sigma Aldrich), *FOXO3* (Mission custom siRNA, V30002, Sigma Aldrich), or negative control scrambled siRNA (Silencer Negative Control No. 1, AM4611, Thermo Fisher). In preparation for transfection, siRNAs in 37.5 μl of OptiMem I Reduced Serum Media (31985, Thermo Fisher) were mixed with an equal volume of OptiMem containing 1.5 μl of Lipofectamine RNAiMax (13778, Thermo Fisher), incubated at room temperature for 15 min, and subsequently added dropwise to cells in 3 ml growth media. Final concentration of siRNAs was 25 nM. In experiments involving stimulated media, an additional round of siRNA transfection was prepared and added dropwise after growth media with decidualizing stimuli was added. Custom, pre-designed siRNAs were acquired in consultation with Sigma-Aldrich for siRNAs targeting *M. domestica FOXO1* and *FOXO3* mRNA. Two siRNAs were transfected for each gene. Custom siRNAs were synthesized by Sigma to target the mRNAs of *M. domestica FOXO1* and *FOXO3,* using the NCBI Reference Sequences XM_001368275.4 (*FOXO1*) and XM_001368456.2 (*FOXO3*). Two custom siRNAs were synthesized for each gene. Sense and antisense sequences are:

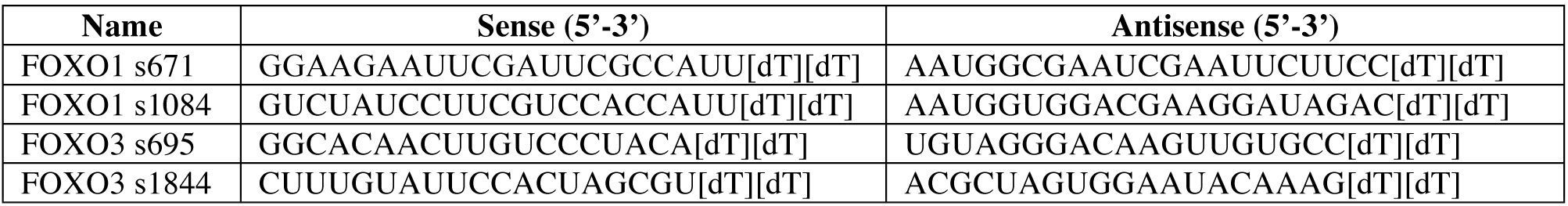

We confirmed depletion of both RNA and protein by qPCR and Western blot. For FOXO1, qPCR analyses showed that knockdown efficiency for these pooled siRNAs were >90% (Extended Data Fig. 4b). This depletion in RNA led to a corresponding depletion in FOXO1 protein, as confirmed by Western blow (Supplementary Information Fig. 3). For FOXO3, we confirmed RNA depletion only (Extended Data Fig. 4b), as we did not find a commercially-available antibody that showed high specificity for FOXO3 in *M. domestica.*

### Flow cytometry including ROS and apoptosis detection

Cells were transfected as above with RNAi reagents, and subsequently incubated in growth media in the presence of decidualizing stimuli for four days. For FACS analyses, conditioned media from each well was decanted into separate 15 ml conical centrifuge tubes, and cells were then washed in 2 ml PBS. PBS was decanted into the same 15 ml conical tube as the conditioned media, and 250 μl warm Tryple Express (12604, Thermo Fisher) was added to each well. To facilitate detachment, cells were placed in a 33°C incubator for 10 min. The detachment reaction was stopped by adding 700 μl growth media to each well. Cells were then transferred to their separate 15 ml conical tubes with conditioned media and PBS, and subsequently pelleted by centrifugation at 400 RPM for 5 min. Supernatant was removed, and each cell pellet was resuspended in 1 ml pre-warmed Hank’s Balanced Salt Solution (HBSS) w/o Phenol Red, Ca^2+^, Mg^2+^ (10-547, Lonza) containing freshly resuspended H_2_DCFDA (Image-IT LIVE Green ROS Detection Kit, I36007, Thermo Fisher) to a final concentration of 25 μM. Cells were incubated at 33°C for 40 min. Just prior to FACS analysis, cells were placed on ice and 0.5 ml HBSS containing μg/ml propidium iodide was added to each tube. We utilized a doublet discrimination gating strategy on a BD Aria FACS instrument, wherein on average 93% of all cells were included in the analyses (Supplementary Information Fig. 9). Fluorescent signal detected in negative control scrambled siRNA cells were utilized to set the quadrant boundaries. Data for three replicates of each experiment were collected and mean percentages for each quadrant were calculated.

### Transcriptomic analyses

Raw sequencing reads were mapped to opossum *M. domestica* genome assembly monDom5 with Ensembl annotation v86, using Tophat2 v2.1.1^54^ and Bowtie2 v2.2.9^55^. Read counts for all genes were calculated using HTSeq v0.6.1p1^56^ with Python (v2.7.13). Transcripts per million were calculated to estimate relative mRNA abundance^57^. We used Bioconductor package edgeR v3.16.5^58^ to assay for differential gene expression between unstimulated and stimulated MdESF. Genes that met the following criteria were considered to be significantly differentially expressed: (1) change in expression of at least 1.5-fold; (2) resulted in an adjusted p-value smaller than 10^−6^; and (3) expressed in at least one condition under comparison (TPM≥3)^59^. Gene ontology enrichment analyses were performed using GOrilla^60^ and the results were visualized using REViGO^61^.

### Statistical analyses

The data from the experiment testing the effect of cAMP/MPA and PGE2/MPA, as well as FOXO1 and FOXO3 knockdowns on the presence of ROS and apoptotic cells, was analyzed as a two factor ANOVA. The two factors were “stimulation” and “treatment”, where stimulation had the levels, “growth media”, cAMP/MPA, and PGE2/MPA, and treatment had the levels random siRNA, FOXO1 KD, and FOXO3 KD. The response variable was the fraction of cells showing either ROS or PI fluoresecence. The analysis was performed with raw frequencies as well a with log-transformed response variables. For the PGE2 ELISA, the concentration of PGE2 for each sample was determined by standard curve. The values were subsequently log transformed and used in a one-tailed t-test.

## Supplementary Information

A supplementary information file is available with this submission. It contains cell line validation data, uncropped Western blot images, unedited REViGO tables for up- and down-regulated genes, uncropped PCR images of gels showing mycoplasma assays, flow cytometry gating analysis, and a table of qPCR primers used in this study.

## Acknowledgements

We appreciate flow cytometry expertise provided by Dr. Ken Nelson of the Yale Center for Molecular Discovery. We thank Dr. Kshitiz for insightful discussions. Transcriptomic data presented in this manuscript were sequenced at the Yale Center for Genome Analysis. This work was funded by John Templeton Foundation Grant #54860 awarded to G.P.W., as well as in part by a Gaylord Donnelley Postdoctoral Environmental Fellowship to O.W.G.

## Author Contributions

E.M.E. designed and performed experiments, analyzed data, wrote and revised the manuscript; J.D.M. performed experiments, analyzed data, revised the manuscript; O.W.G. performed experiments, analyzed data, wrote and revised the manuscript; C.L. analyzed data; A.R.C. designed experiments, revised the manuscript; G.P.W. conceived the study, designed experiments, analyzed data, wrote and revised the manuscript.

## Data availability

All RNAseq data in this study has been uploaded to GEO (accession number available upon acceptance).

## Extended Data

**Extended Data Table 1.** List of 28 transcription factors with documented roles in decidualization with references.

| Transcription Factor | Reference(s) |
| --- | --- |
| *CEBPB* | 17,20 |
| *E2F8* | 41 |
| *EBF4* | 7 |
| *ETS1* | 62 |
| *FOS* | 63 |
| *FOXM1* | 64 |
| *FOXO1* | 18 |
| *GATA2* | 25 |
| *HAND2* | 31 |
| *HOXA10* | 21,22 |
| *HOXA11* | 23,24 |
| *HOXD9* | 16 |
| *HOXD11* | 65 |
| *HOXD12* | 16 |
| *KLF9* | 28-30 |
| *MEIS1* | 34 |
| *NR2F2* | 66 |
| *PBX2* | 67 |
| *PGR* | 19 |
| *PRRX2* | 68 |
| *RUNX1* | 69 |
| *STAT3* | 32,33 |
| *STAT5B* | 70 |
| *TFAP2C* | 16 |
| *TWIST1* | 71 |
| *WT1* | 72 |
| *ZBTB16* | 26,27 |

## Extended Data Figures

**Extended Data Figure 1.**
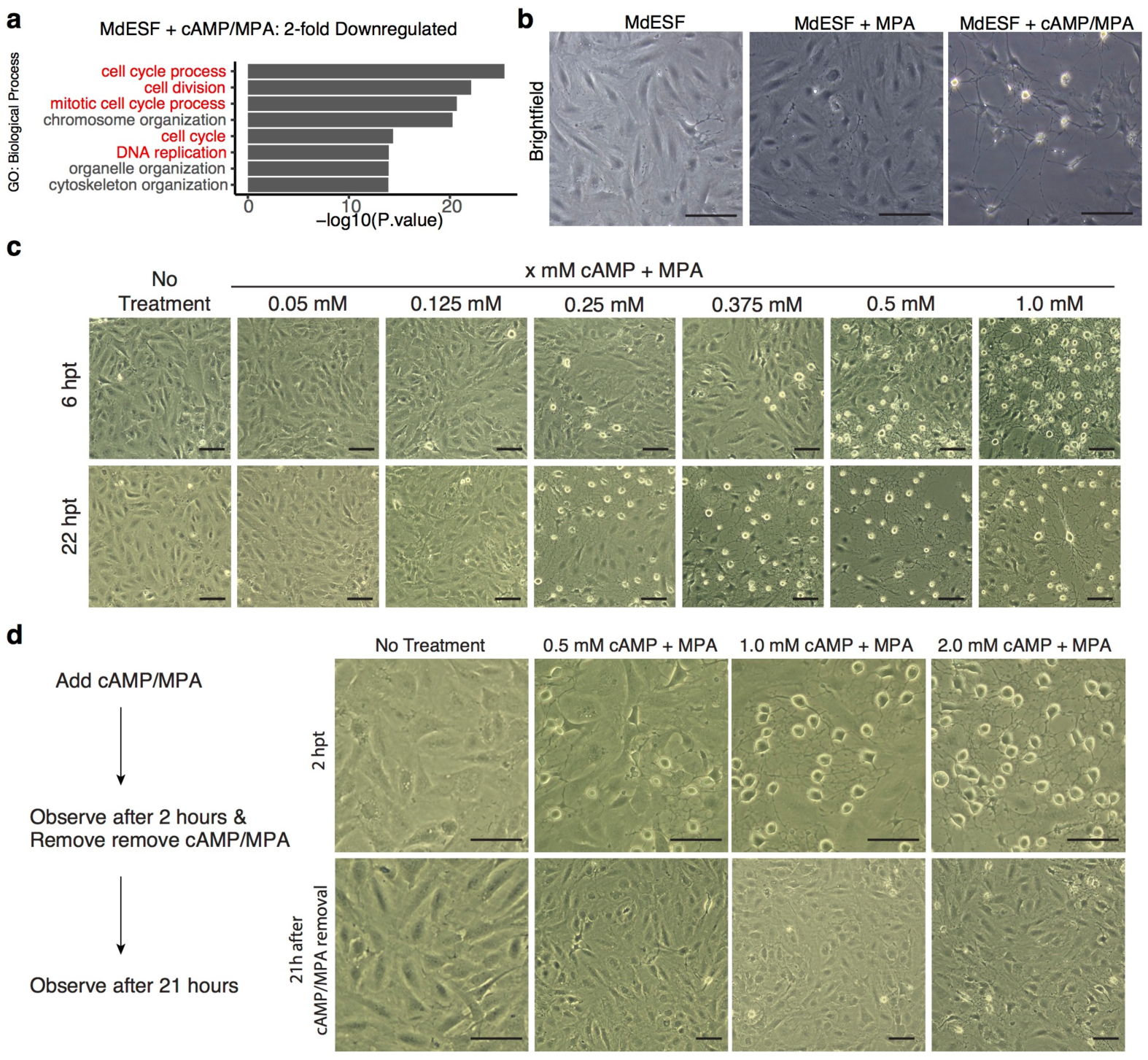
Transcriptional, cAMP-concentration dependent, and reversible response of MdESF to decidualizing stimuli. **a,** GO categories of downregulated genes in MdESF after treatment with decidualizing stimuli relative to unstimulated control (n=3, p-value<10^−6^). **b,** DIC images showing morphological response of MdESF treated with either MPA or cAMP/MPA. Images obtained during ROS detection (see Fig. 1d). **c,** Concentration-dependent morphological response of MdESF to cAMP/MPA treatment after six hours (top panels) and 22 hours (bottom panels). **d,** The MdESF morphological response to cAMP/MPA treatment is reversible. Decidualizing stimuli were added at the three different concentrations shown. Images were taken two hours after treatment, at which time media were changed to growth media. Images were subsequently acquired next day after 19 hours in growth media. Scale bars are 10 μM.

**Extended Data Figure 2.**
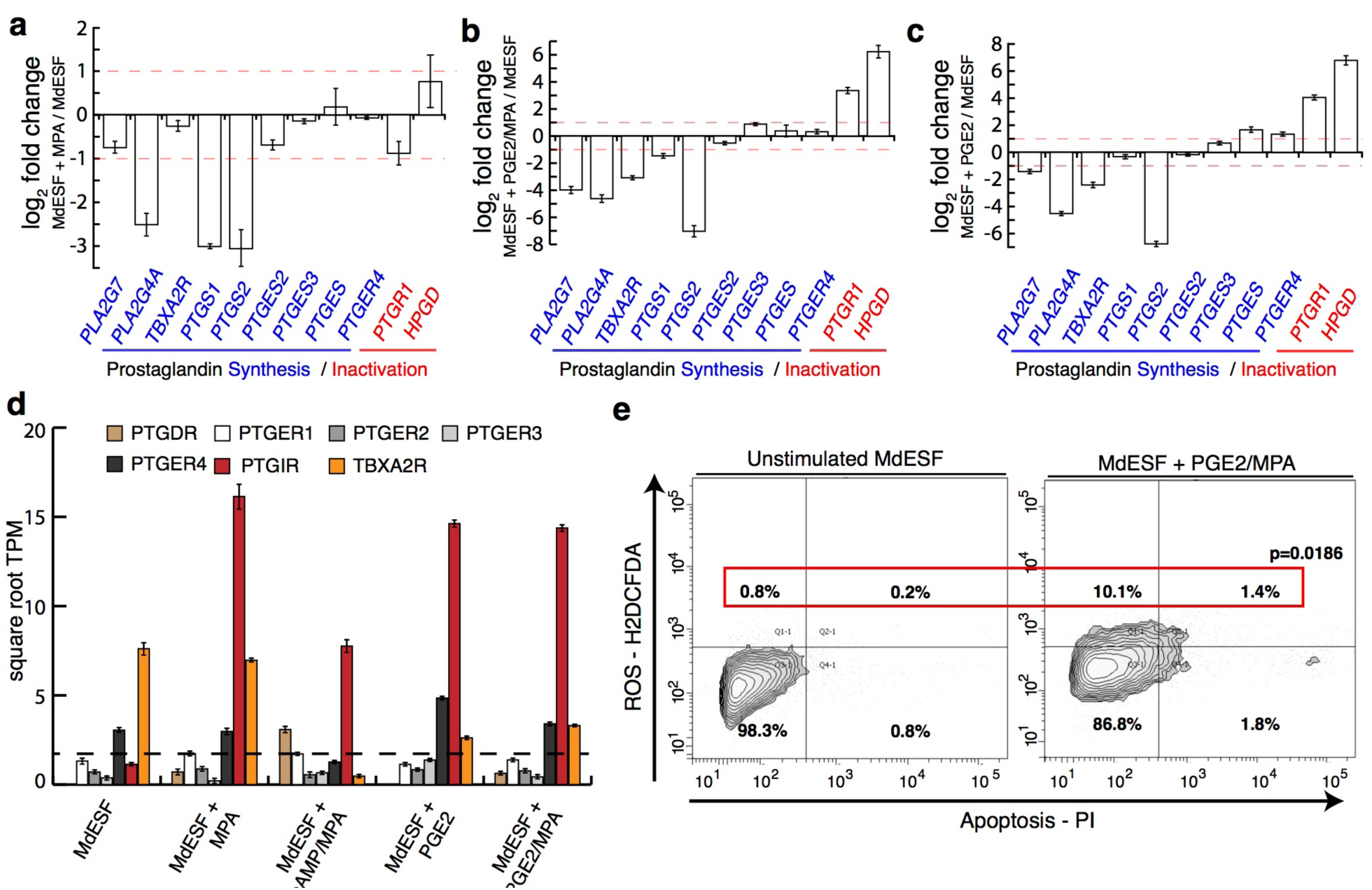
Transcriptional response of KEGG pathway genes associated with prostaglandin signaling in MdESF treated with stimuli in this study. **a-c,** Transcriptional response of prostaglandin synthesis and inactivation genes in MdESF treated with **a,** MPA alone, **b,** PGE2 alone, or **c,** PGE2/MPA. Fold change relative to unstimulated control is shown (n=3, log2 fold change shown). Red dashed line represents 2-fold change. **d,** Expression of orthologous receptors in MdESF for prostaglandin signaling pathways in unstimulated MdESF or stimulated with MPA alone, cAMP/MPA, PGE2 alone, or PGE2/MPA. Colors represent receptor indicated in legend. Average square root transcripts per million (TPM) of three replicates is shown. Error bars show standard deviation of the mean. **e,** Flow cytometry contour plots showing ROS (H2DCFDA) versus staining for apoptosis (propidium iodide). Quadrants are set to maximum extent of boundaries in unstimulated MdESF treated with scrambled siRNA. Q1, ROS positive only; Q2, ROS and apoptosis positive; Q3, negative for ROS and apoptosis; Q4, negative for ROS and positive for apoptosis. Percentage in each quadrant for this replicate is shown. Note increase of ROS positive cells after PGE2/MPA treatment, similar to what was found with cAMP/MPA (p=0.0186).

**Extended Data Figure 3.**
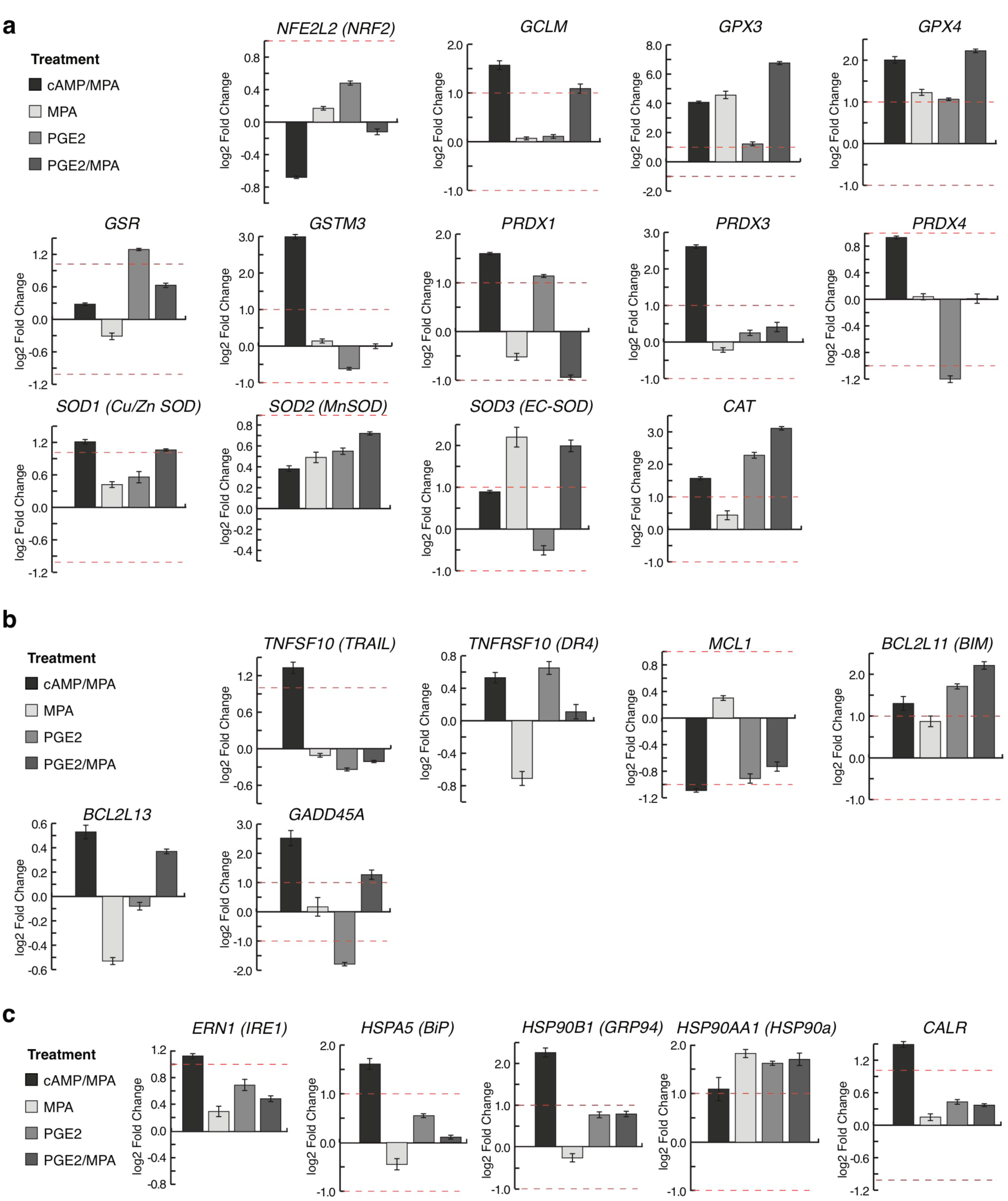
Stress-related gene expression changes in all MdESF treatments relative to unstimulated control. Genes of interest are shown that are known to be directly involved at the transcriptional level in **a,** oxidative stress response; **b,** apoptosis; **c,** unfolded protein response in the endoplasmic reticulum due to oxidative stress.

**Extended Data Figure 4.**
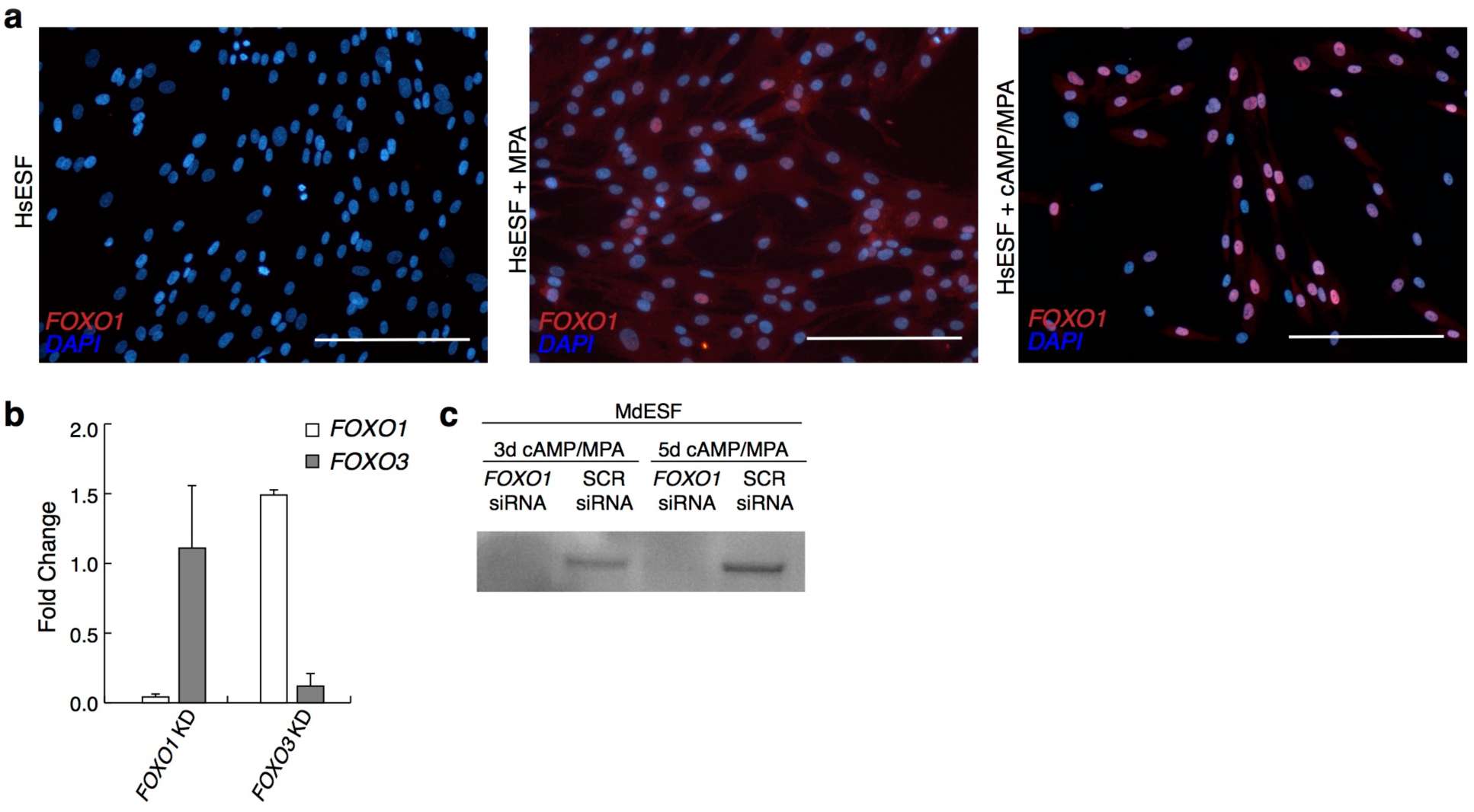
Immunofluorescence of FOXO1 in HsESF and FOXO1/FOXO3 knockdown in MdESF. **a,** Although *FOXO1* RNA is present in HsESF, in HsESF it is constantly marked for degradation by polyubiquitination by AKT. In the presence of MPA, degradation of FOXO1 protein is disrupte **d,** and FOXO1 disproportionately loads in the cytoplasm relative to the nucleus, though some cells are positive for nuclear FOXO1. In the presence of cAMP/MPA, FOXO1 protein loads disproportionately in the nucleus relative to the cytoplasm in HsESF. Scale bars are 20 μM. **b,** Fold change of *FOXO1* and *FOXO3* RNA in cells treated with siRNA targeting *FOXO1* and *FOXO3* relative to scrambled siRNA control. siRNAs targeting *FOXO1* and *FOXO3* RNA removed greater than ≥ 90% of *FOXO1* and *FOXO3* transcripts. **c,** Western blot for FOXO1 in total protein lysates collected from MdESF treated with cAMP/MPA for 3 days or 5 days and with siRNA targeting *FOXO1* RNA.

## References

1 Liang, C., Consortium, F., Forrest, A. R. & Wagner, G. P. The statistical geometry of transcriptome divergence in cell-type evolution and cancer. Nat Commun 6, 6066, doi:10.1038/ncomms7066 (2015).

2 Arendt, D. The evolution of cell types in animals: emerging principles from molecular studies. Nat Rev Genet 9, 868–882, doi:10.1038/nrg2416 (2008).

3 Arendt, D. et al. The origin and evolution of cell types. Nat Rev Genet 17, 744–757, doi:10.1038/nrg.2016.127 (2016).

4 Griffith, O. W. & Wagner, G. P. The placenta as a model for understanding the origin and evolution of vertebrate organs. Nat Ecol Evol 1, 72, doi:10.1038/s41559-017-0072 (2017).

5 Chavan, A. R., Griffith, O. W. & Wagner, G. P. The inflammation paradox in the evolution of mammalian pregnancy: turning a foe into a friend. Curr Opin Genet Dev 47, 24–32, doi:10.1016/j.gde.2017.08.004 (2017).

6 Gellersen, B. & Brosens, J. Cyclic AMP and progesterone receptor cross-talk in human endometrium: a decidualizing affair. J Endocrinol 178, 357–372 (2003).

7 Kin, K., Nnamani, M. C., Lynch, V. J., Michaelides, E. & Wagner, G. P. Cell-type phylogenetics and the origin of endometrial stromal cells. Cell Rep 10, 1398–1409, doi:10.1016/j.celrep.2015.01.062 (2015).

8 Mess, A. & Carter, A. M. Evolutionary transformations of fetal membrane characters in Eutheria with special reference to Afrotheria. J Exp Zool B Mol Dev Evol 306, 140–163, doi:10.1002/jez.b.21079 (2006).

9 O’Leary, M. A. et al. The placental mammal ancestor and the post-K-Pg radiation of placentals. Science 339, 662–667, doi:10.1126/science.1229237 (2013).

10 Luo, Z. X., Yuan, C. X., Meng, Q. J. & Ji, Q. A Jurassic eutherian mammal and divergence of marsupials and placentals. Nature 476, 442–445, doi:10.1038/nature10291 (2011).

11 Frank, G. R., Brar, A. K., Cedars, M. I. & Handwerger, S. Prostaglandin E2 enhances human endometrial stromal cell differentiation. Endocrinology 134, 258–263, doi:10.1210/endo.134.1.7506205 (1994).

12 Fortier, M. A., Boulanger, M., Boulet, A. P. & Lambert, R. D. Cell-specific localization of prostaglandin E2-sensitive adenylate cyclase in rabbit endometrium. Biol Reprod 36, 1025–1033 (1987).

13 Yee, G. M. & Kennedy, T. G. Role of cyclic adenosine 3’,5’-monophosphate in mediating the effect of prostaglandin E2 on decidualization in vitro. Biol Reprod 45, 163–171 (1991).

14 Lane, B., Oxberry, W., Mazella, J. & Tseng, L. Decidualization of human endometrial stromal cells in vitro: effects of progestin and relaxin on the ultrastructure and production of decidual secretory proteins. Hum Reprod 9, 259–266 (1994).

15 Gellersen, B. & Brosens, J. J. Cyclic decidualization of the human endometrium in reproductive health and failure. Endocr Rev 35, 851–905, doi:10.1210/er.2014-1045 (2014).

16 Kin, K. et al. The transcriptomic evolution of mammalian pregnancy: Gene expression innovations in endometrial stromal fibroblasts. Genome Biol Evol 8, 2459–2473, doi:10.1093/gbe/evw168 (2016).

17 Christian, M. et al. Cyclic AMP-induced forkhead transcription factor, FKHR, cooperates with CCAAT/enhancer-binding protein beta in differentiating human endometrial stromal cells. J Biol Chem 277, 20825–20832, doi:10.1074/jbc.M201018200 (2002).

18 Grinius, L., Kessler, C., Schroeder, J. & Handwerger, S. Forkhead transcription factor FOXO1A is critical for induction of human decidualization. J Endocrinol 189, 179–187, doi:10.1677/joe.1.06451 (2006).

19 Tseng, L. & Zhu, H. H. Regulation of progesterone receptor messenger ribonucleic acid by progestin in human endometrial stromal cells. Biol Reprod 57, 1360–1366 (1997).

20 Mantena, S. R. et al. C/EBPbeta is a critical mediator of steroid hormone-regulated cell proliferation and differentiation in the uterine epithelium and stroma. Proc Natl Acad Sci U S A 103, 1870–1875, doi:10.1073/pnas.0507261103 (2006).

21 Taylor, H. S., Arici, A., Olive, D. & Igarashi, P. HOXA10 is expressed in response to sex steroids at the time of implantation in the human endometrium. J Clin Invest 101, 1379–1384, doi:10.1172/JCI1057 (1998).

22 Godbole, G. & Modi, D. Regulation of decidualization, interleukin-11 and interleukin-15 by homeobox A 10 in endometrial stromal cells. J Reprod Immunol 85, 130–139, doi:10.1016/j.jri.2010.03.003 (2010).

23 Taylor, H. S., Igarashi, P., Olive, D. L. & Arici, A. Sex steroids mediate HOXA11 expression in the human peri-implantation endometrium. J Clin Endocrinol Metab 84, 1129–1135, doi:10.1210/jcem.84.3.5573 (1999).

24 Gendron, R. L. et al. Abnormal uterine stromal and glandular function associated with maternal reproductive defects in Hoxa-11 null mice. Biol Reprod 56, 1097–1105 (1997).

25 Rubel, C. A. et al. Gata2 is a master regulator of endometrial function and progesterone signaling. Biol Reprod 85, 179 (2011).

26 Kommagani, R. et al. The promyelocytic leukemia zinc finger transcription factor is critical for human endometrial stromal cell decidualization. PLoS Genet 12, e1005937, doi:10.1371/journal.pgen.1005937 (2016).

27 Szwarc, M. M. et al. Human endometrial stromal cell decidualization requires transcriptional reprogramming by PLZF. Biol Reprod, doi:10.1093/biolre/iox161 (2017).

28 Pabona, J. M., Zeng, Z., Simmen, F. A. & Simmen, R. C. Functional differentiation of uterine stromal cells involves cross-regulation between bone morphogenetic protein 2 and Kruppel-like factor (KLF) family members KLF9 and KLF13. Endocrinology 151, 3396–3406, doi:10.1210/en.2009-1370 (2010).

29 Simmen, R. C. et al. Subfertility, uterine hypoplasia, and partial progesterone resistance in mice lacking the Kruppel-like factor 9/basic transcription element-binding protein-1 (Bteb1) gene. J Biol Chem 279, 29286–29294, doi:10.1074/jbc.M403139200 (2004).

30 Pabona, J. M. et al. Kruppel-like factor 9 and progesterone receptor coregulation of decidualizing endometrial stromal cells: implications for the pathogenesis of endometriosis. J Clin Endocrinol Metab 97, E376–392, doi:10.1210/jc.2011-2562 (2012).

31 Li, Q. et al. The antiproliferative action of progesterone in uterine epithelium is mediated by Hand2. Science 331, 912–916, doi:10.1126/science.1197454 (2011).

32 Dimitriadis, E., Stoikos, C., Tan, Y. L. & Salamonsen, L. A. Interleukin 11 signaling components signal transducer and activator of transcription 3 (STAT3) and suppressor of cytokine signaling 3 (SOCS3) regulate human endometrial stromal cell differentiation. Endocrinology 147, 3809–3817, doi:10.1210/en.2006-0264 (2006).

33 Wang, W., Taylor, R. N., Bagchi, I. C. & Bagchi, M. K. Regulation of human endometrial stromal proliferation and differentiation by C/EBPbeta involves cyclin E-cdk2 and STAT3. Mol Endocrinol 26, 2016–2030, doi:10.1210/me.2012-1169 (2012).

34 Hu, L. et al. Regulation of myeloid ecotropic viral integration site 1 and its expression in normal and abnormal endometrium. Fertil Steril 102, 856–863 e852, doi:10.1016/j.fertnstert.2014.05.036 (2014).

35 Griffith, O. W. et al. Embryo implantation evolved from an ancestral inflammatory attachment reaction. Proc Natl Acad Sci U S A 114, E6566–E6575, doi:10.1073/pnas.1701129114 (2017).

36 Griffith, O. W. et al. Reply to Liu: Inflammation before implantation both in evolution and development. Proc Natl Acad Sci U S A 115, E3–E4, doi:10.1073/pnas.1717001115 (2018).

37 Huang, H. & Tindall, D. J. Dynamic FoxO transcription factors. J Cell Sci 120, 2479–2487, doi:10.1242/jcs.001222 (2007).

38 Kajihara, T., Brosens, J. J. & Ishihara, O. The role of FOXO1 in the decidual transformation of the endometrium and early pregnancy. Med Mol Morphol 46, 61–68, doi:10.1007/s00795-013-0018-z (2013).

39 Kin, K., Maziarz, J. & Wagner, G. P. Immunohistological study of the endometrial stromal fibroblasts in the opossum, Monodelphis domestica: evidence for homology with eutherian stromal fibroblasts. Biol Reprod 90, 111, doi:10.1095/biolreprod.113.115139 (2014).

40 Bridge, D. et al. FoxO and stress responses in the cnidarian Hydra vulgaris. PLoS One 5, e11686, doi:10.1371/journal.pone.0011686 (2010).

41 Qi, Q. R. et al. Involvement of atypical transcription factor E2F8 in the polyploidization during mouse and human decidualization. Cell Cycle 14, 1842–1858, doi:10.1080/15384101.2015.1033593 (2015).

42 Hawkins, M. & Battaglia, A. Breeding behaviour of the platypus (Ornithorhynchus anatinus) in captivity. Aust J Zool 57, 283–293 (2009).

43 Hall, B. K. The paradoxical platypus. BioScience 49, 211–218 (1999).

44 Kajihara, T. et al. Differential expression of FOXO1 and FOXO3a confers resistance to oxidative cell death upon endometrial decidualization. Mol Endocrinol 20, 2444–2455, doi:10.1210/me.2006-0118 (2006).

45 Al-Sabbagh, M. et al. NADPH oxidase-derived reactive oxygen species mediate decidualization of human endometrial stromal cells in response to cyclic AMP signaling. Endocrinology 152, 730–740, doi:10.1210/en.2010-0899 (2011).

46 Maruyama, T. et al. Induction of thioredoxin, a redox-active protein, by ovarian steroid hormones during growth and differentiation of endometrial stromal cells in vitro. Endocrinology 140, 365–372, doi:10.1210/endo.140.1.6455 (1999).

47 Maruyama, T. et al. Thioredoxin expression in the human endometrium during the menstrual cycle. Mol Hum Reprod 3, 989–993 (1997).

48 Labied, S. et al. Progestins regulate the expression and activity of the forkhead transcription factor FOXO1 in differentiating human endometrium. Mol Endocrinol 20, 35–44, doi:10.1210/me.2005-0275 (2006).

49 Guzel, E. et al. Endoplasmic reticulum stress and homeostasis in reproductive physiology and pathology. Int J Mol Sci 18, doi:10.3390/ijms18040792 (2017).

50 Brighton, P. J. et al. Clearance of senescent decidual cells by uterine natural killer cells in cycling human endometrium. Elife 6, doi:10.7554/eLife.31274 (2017).

51 Borquez, D. A. et al. Dissecting the role of redox signaling in neuronal development. J Neurochem 137, 506–517, doi:10.1111/jnc.13581 (2016).

52 Wilson, C., Munoz-Palma, E. & Gonzalez-Billault, C. From birth to death: A role for reactive oxygen species in neuronal development. Semin Cell Dev Biol, doi:10.1016/j.semcdb.2017.09.012 (2017).

53 Denu, R. A. & Hematti, P. Effects of oxidative stress on mesenchymal stem cell biology. Oxid Med Cell Longev 2016, 2989076, doi:10.1155/2016/2989076 (2016).

54 Kim, D. et al. TopHat2: accurate alignment of transcriptomes in the presence of insertions, deletions and gene fusions. Genome Biol 14, R36, doi:10.1186/gb-2013-14-4-r36 (2013).

55 Langmead, B. & Salzberg, S. L. Fast gapped-read alignment with Bowtie 2. Nat Methods 9, 357–359, doi:10.1038/nmeth.1923 (2012).

56 Anders, S., Pyl, P. T. & Huber, W. HTSeq‐‐a Python framework to work with high-throughput sequencing data. Bioinformatics 31, 166–169, doi:10.1093/bioinformatics/btu638 (2015).

57 Wagner, G. P., Kin, K. & Lynch, V. J. Measurement of mRNA abundance using RNA-seq data: RPKM measure is inconsistent among samples. Theory Biosci 131, 281–285, doi:10.1007/s12064-012-0162-3 (2012).

58 Robinson, M. D., McCarthy, D. J. & Smyth, G. K. edgeR: a Bioconductor package for differential expression analysis of digital gene expression data. Bioinformatics 26, 139–140, doi:10.1093/bioinformatics/btp616 (2010).

59 Wagner, G. P., Kin, K. & Lynch, V. J. A model based criterion for gene expression calls using RNA-seq data. Theory Biosci 132, 159–164, doi:10.1007/s12064-013-0178-3 (2013).

60 Eden, E., Navon, R., Steinfeld, I., Lipson, D. & Yakhini, Z. GOrilla: a tool for discovery and visualization of enriched GO terms in ranked gene lists. BMC Bioinformatics 10, 48, doi:10.1186/1471-2105-10-48 (2009).

61 Supek, F., Bosnjak, M., Skunca, N. & Smuc, T. REVIGO summarizes and visualizes long lists of gene ontology terms. PLoS One 6, e21800, doi:10.1371/journal.pone.0021800 (2011).

62 Kessler, C. A., Schroeder, J. K., Brar, A. K. & Handwerger, S. Transcription factor ETS1 is critical for human uterine decidualization. Mol Hum Reprod 12, 71–76, doi:10.1093/molehr/gal008 (2006).

63 Reis, F. M., Maia, A. L., Ribeiro, M. F. & Spritzer, P. M. Progestin modulation of c-fos and prolactin gene expression in the human endometrium. Fertil Steril 71, 1125–1132 (1999).

64 Gao, F., Bian, F., Ma, X., Kalinichenko, V. V. & Das, S. K. Control of regional decidualization in implantation: Role of FoxM1 downstream of Hoxa10 and cyclin D3. Sci Rep 5, 13863, doi:10.1038/srep13863 (2015).

65 Akbas, G. E. & Taylor, H. S. HOXC and HOXD gene expression in human endometrium: lack of redundancy with HOXA paralogs. Biol Reprod 70, 39–45, doi:10.1095/biolreprod.102.014969 (2004).

66 Li, X. et al. COUP-TFII regulates human endometrial stromal genes involved in inflammation. Mol Endocrinol 27, 2041–2054, doi:10.1210/me.2013-1191 (2013).

67 Sarno, J. L., Kliman, H. J. & Taylor, H. S. HOXA10, Pbx2, and Meis1 protein expression in the human endometrium: formation of multimeric complexes on HOXA10 target genes. J Clin Endocrinol Metab 90, 522–528, doi:10.1210/jc.2004-0817 (2005).

68 Jiang, Y. et al. Uterine Prx2 restrains decidual differentiation through inhibiting lipolysis in mice. Cell Tissue Res 365, 403–414, doi:10.1007/s00441-016-2383-0 (2016).

69 Bai, Z. K. et al. Differential expression and regulation of Runx1 in mouse uterus during the peri-implantation period. Cell Tissue Res 362, 231–240, doi:10.1007/s00441-015-2174-z (2015).

70 Mak, I. Y. et al. Regulated expression of signal transducer and activator of transcription, Stat5, and its enhancement of PRL expression in human endometrial stromal cells in vitro. J Clin Endocrinol Metab 87, 2581–2588, doi:10.1210/jcem.87.6.8576 (2002).

71 Schroeder, J. K., Kessler, C. A. & Handwerger, S. Critical role for TWIST1 in the induction of human uterine decidualization. Endocrinology 152, 4368–4376, doi:10.1210/en.2011-1140 (2011).

72 Tamura, I. et al. Novel Function of a Transcription Factor WT1 in Regulating Decidualization in Human Endometrial Stromal Cells and Its Molecular Mechanism. Endocrinology 158, 3696–3707, doi:10.1210/en.2017-00478 (2017).

